# Structural Flexibility and Disassembly Kinetics of Single Ferritins Using Optical Nanotweezers

**DOI:** 10.1101/2023.09.22.558948

**Authors:** Arman Yousefi, Ze Zheng, Saaman Zargarbashi, Mahya Assadipapari, Graham J. Hickman, Christopher D. J. Parmenter, Carlos J. Bueno-Alejo, Gabriel Sanderson, Dominic Craske, Lei Xu, Mohsen Rahmani, Cuifeng Ying

## Abstract

Ferritin, a spherical protein shell assembled from 24 subunits, functions as an efficient iron storage and release system through its channels. Understanding how various chemicals affect the structural behaviour of ferritin is crucial for unravelling the origins of iron-related diseases in living organisms including humans. In particular, the influence of chemicals on ferritin’s dynamics and iron release is barely explored at the single-protein level. Here, by employing optical nanotweezers using double nanohole (DNH) structures, we examined the effect of ascorbic acid (reducing reagent) and pH on ferritin’s conformational dynamics. The dynamics of ferritin increased as the concentration of ascorbic acid approached saturation. At pH 2.0 ferritin exhibited significant structural fluctuations and eventually underwent a stepwise disassembly into fragments. This work, for the first time, tracked the disassembly pathway and kinetics of single ferritins in solution. We identified four critical fragments during its disassembly pathway, which are 22-mer, 12-mer, tetramer, and dimer subunits. Moreover, we presented the first single-molecule evidence of the cooperative disassembly of ferritin. Interrogating ferritin’s structural change in response to different chemicals holds importance for understanding their roles in iron metabolism, hence facilitating further development of medical treatments for the associated diseases.

---

Ferritin is a primary iron storage protein, consisting of a spherical cage formed by 24 identical subunits (24-mer) arranged in 3-fold and 4-fold channels on the shell.^1,2^ Ferritin sequesters iron within a ferrihydrite (Fe^3+^) core and releases it as ferrous iron (Fe^2+^) in response to biological demand. Ferrous iron plays a crucial role in preventing oxidative damage to cells through its participation in the Fenton reaction, which involves the decomposition of hydrogen peroxide and its interaction with reactive oxygen species.^3,4^ Various reductants can facilitate iron mobilisation from the ferritin shell through 3-fold channels.^5,6^ For instance, ascorbic acid (AA) facilitates cellular metabolism and serves as a reducing agent for mobilising iron in ferritin.^7^ Patients with hemochromatosis, a disorder characterised by significant iron overload, have lower ascorbate levels than normal and, therefore, require supplementation of ascorbate during chelation therapy.^8,9^ High ascorbate levels can cause damage to proteins, lipids, and DNA by generating radicals via the Fenton reaction.^10^ Additionally, high ascorbate concentrations can create an acidic environment, potentially impacting ferritin dynamics and iron release,^11^ as ferritin is known to undergo disassembly at extreme acid conditions (*i.e.*, pH ≤ 2). The disassembled ferritin can reassemble at higher pH due to electrostatic interactions.^12^ These features make ferritin promising for biomedical applications like drug delivery^13,14^ and imaging,^15,16^ However, a deep understanding of its behaviour in response to different conditions is essential for effective utilisation in these areas.^17^

Various analytical approaches, including gel electrophoresis,^18^ circular dichroism,^19^ ultracentrifugation,^20^ and fluorescence microscopy,^21^ have revealed the reversible disassembly and assembly of ferritin in response to pH changes. The pH-dependent structural alterations and assembly kinetics of ferritin have also been demonstrated in detail using small-angle X-ray scattering (SAXS).^12^ These bulk measurements, however, are limited to providing information about an average response for a protein population.^22^ Recently, using high-speed atomic force microscopy (HS-AFM) and molecular dynamic (MD) simulations, Maity *et al.* ^23^ observed real-time disassembly-reassembly of single ferritins at acidic pH conditions. They revealed two disassembly stages: pore formation through the 3-fold channels and subsequent fragmentation into dimers. This study offers valuable insights into subunit-subunit interactions during protein disassembly. However, certain intermediate steps and small domain movements in the disassembly process are not fully captured due to the high stiffness of the AFM cantilever.^24^ Molecular Dynamic simulation, on the other hand, is restricted by its short simulation time (*i.e*., ∼nanoseconds), making it difficult to track large domain movements of biological entities accurately. Therefore, a comprehensive understanding of the dynamic and kinetic behaviour of single, native ferritins, which link to iron binding, releasing and interaction with other biomolecules, remains challenging using these methods.

Using the plasmonic optical tweezers, our recent work demonstrated the ability to monitor the dynamic changes of individual, unmodified ferritins during *in situ* iron loading.^25^ In this study, we investigated the structural flexibility and disassembly kinetics of single ferritins when exposed to ascorbic acid (reducing agent) and acidic environment. We show that ferritin exhibits increased structural fluctuations at ascorbic acid concentrations approaching saturation, attributed to the large amount of iron release. Additionally, when exposing ferritin to acidic conditions at pH 2, the protein became unstable and underwent stepwise disassembly. For the first time, we tracked the disassembly pathway of single ferritins and the time duration of each intermediate fragment during the disassembly. Furthermore, using a recent application of interferometric scattering called Mass Photometry (MP),^26^ we confirmed the existence of these intermediate fragments in ferritin at pH 2. Understanding the disassembly process of ferritin could contribute to the crucial initial step in utilising ferritin for nanotechnology-based applications such as drug delivery and bio-imaging.

## Results and Discussion

### Ascorbic acid opens pore channels of ferritin

Figure 1a depicts the DNH structure in an aqueous environment within an optical nanotweezer setup, where an 852 nm laser is focused on the structure. The localised surface plasmonic resonance generated by the gold DNH highly enhanced the optical field, as shown by the simulated electric field distribution in Figure 1b, with detailed simulation parameters in SI-2. Subsequently, the tightly confined optical field generates a strong gradient force that enables trapping a nanoparticle into the gap of the DNH.^27–31^ The presence of the protein in the gap introduces a resonance shift, leading to a change in the transmission detected by the Avalanche Photodiode (APD). The top panel of Figure 1b demonstrates a typical trapping trace, where the transmission increases upon trapping a single ferritin due to the dielectric loading. ^32–35^

**Figure 1.**
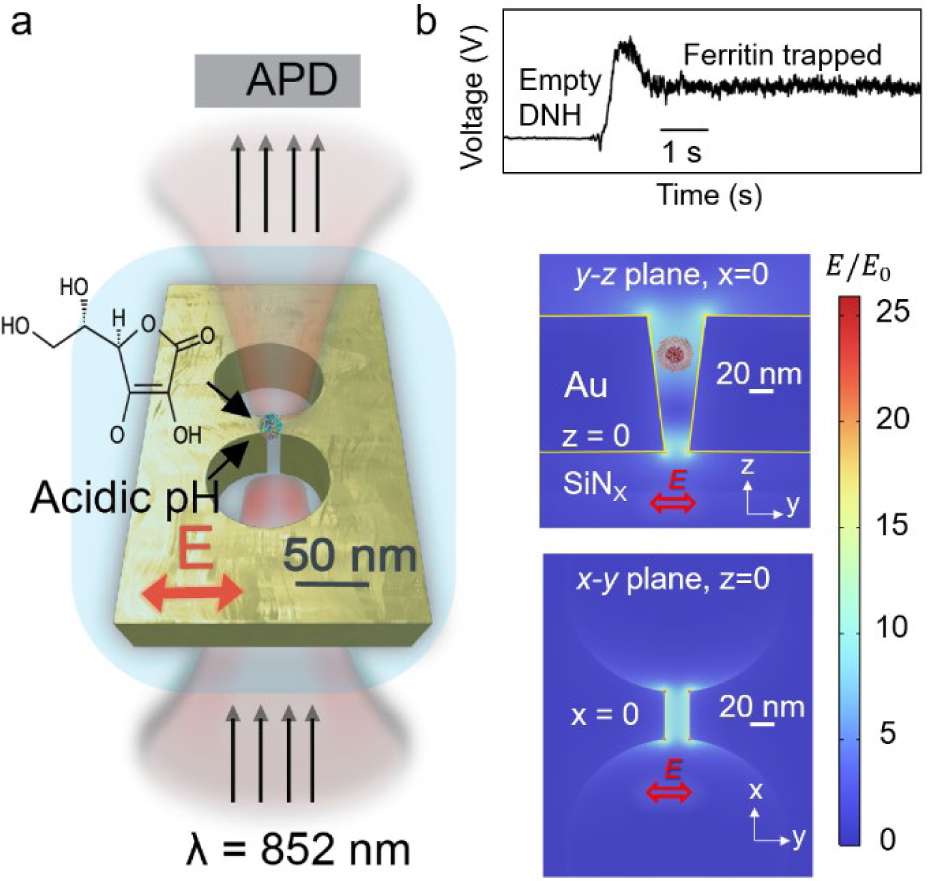
Concept of DNH-based optical nanotweezer with the simulated optical field distribution. (a) Schematic representation of a gold DNH positioned within the flow cell (light blue), with an 852 nm laser beam focused on the structure. Solutions of varying pHs and different concentrations of ascorbic acid are introduced to the trapping site. The transmitted light through the DNH is recorded by an avalanche photodiode (APD). (b) Top panel: representative transmission trace demonstrating the trapping of a single ferritin. Middle and bottom panels: simulated distribution of electric field enhancement in the DNH structure in the y-z plane and x-y plane, respectively.

Ascorbic acid reduces ferric iron into ferrous iron, a process that results in an increased production of reactive oxygen species, specifically hydroxyl radicals.^36^ Subsequently, this oxidation process gives rise to the formation of ascorbyl radicals and ferrous ions (Eq. 1). The ascorbyl radical later reacts with dioxygen, leading to the regeneration of ascorbate and the generation of superoxide ion (Eq. 2). Both the superoxide and ascorbate radicals interact with ferritin, facilitating the conversion of ferric iron into ferrous iron and thereby promoting the mobilisation of iron (Eqs. 3 and 4).^37,38^

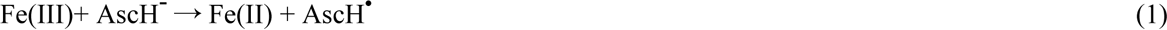

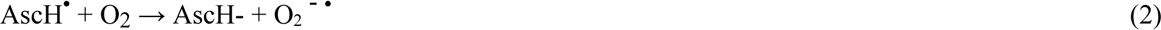

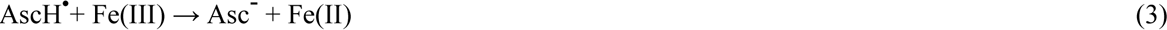

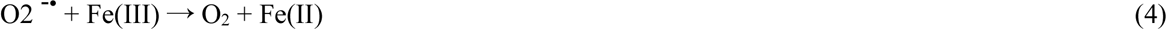

Figure 2a illustrates the process of ascorbate diffusion into the ferritin core through the 3-fold channels,^39^ binding to the ferric ions (Fe^3+^) at the site of the ferroxidase centre, leading to the ferric ions reduction and eventually released from the protein. Figure 2b displays the transmission signals of the trapped ferritin when subjected to varying concentrations of ascorbic acid. As the transmitted light intensity correlates to the conformation of the trapped protein, the root-mean-square (RMS) of the optical signals reveals the fluctuation of protein conformations.^25^ The RMS of the traces in Figure 2b demonstrates an increased dynamic of ferritin within high concentrations of ascorbic acid, arising from the opening and closing of the 3-fold channels due to the permeation of ascorbate and ferrous iron. The pro-oxidant behaviour of high concentrations of ascorbic acid promotes the protein’s channels opening and induces increased amplitude of the transmission signal. Additionally, the loss of the ferric core after exposure to different ascorbate concentrations may contribute to the enhanced structural instability.

**Figure 2.**
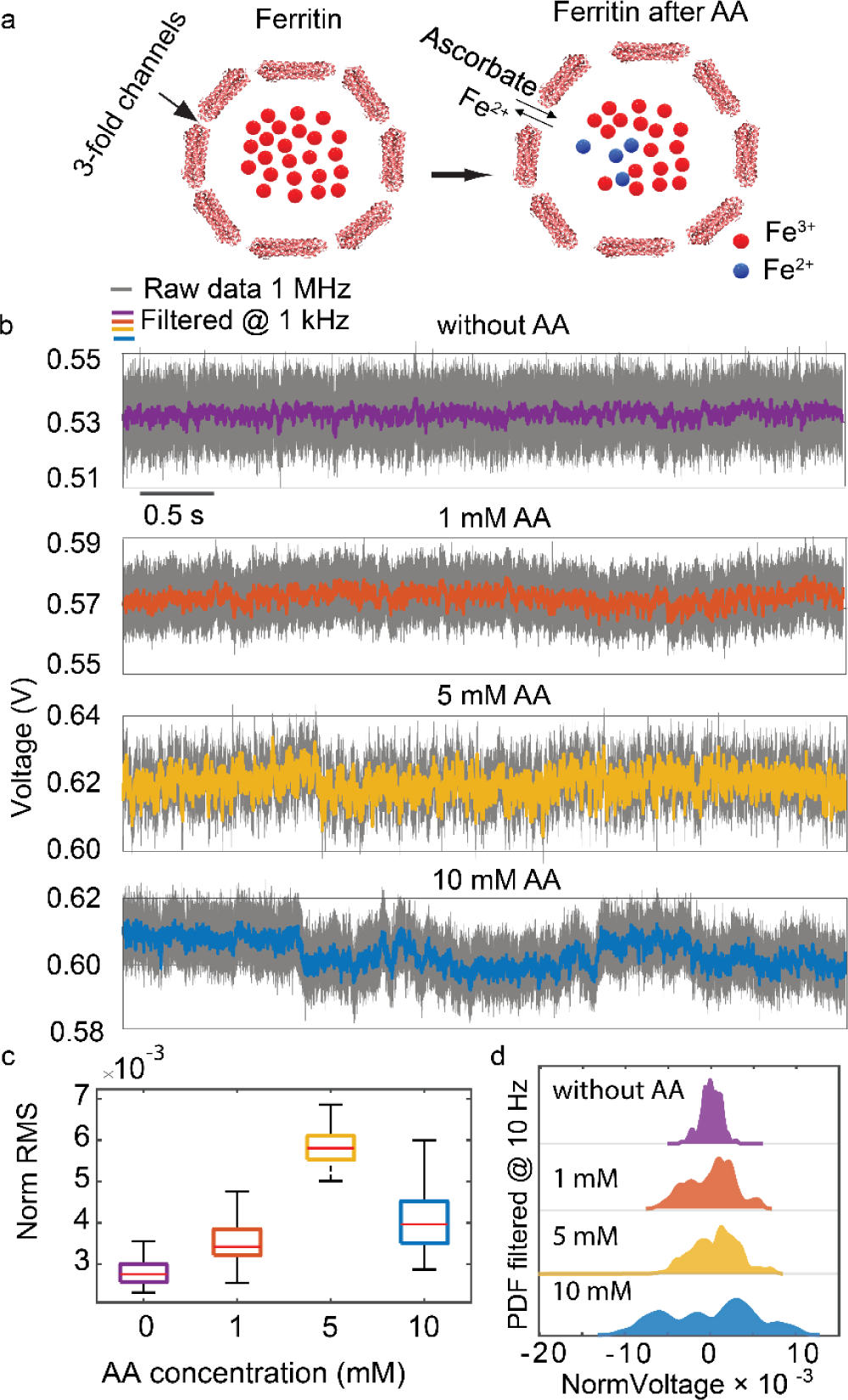
Effect of the ascorbic acid on the dynamic and iron releasing of ferritin. (a) The cartoon illustrates ferritin before and after exposure to ascorbic acid and how ascorbate and Fe2+ permeate through 3-fold channels. (b) Transmission signal of DNH with trapped ferritin within different concentrations of ascorbic acid (0 mM, 1 mM, 5 mM, and 10 mM). (c) Normalised RMS of 5-second transmission signal when ferritin is trapped and introduced to different ascorbic acid concentrations. The data were Gaussian-filtered at 1 kHz, and the box plot displays the 25th and 75th percentiles of the data(d) Probability density function (PDF) of 5-second traces of single ferritin and different ascorbic acid concentrations (filtered at 10 Hz). Data were acquired at 1 MHz and then digitally filtered at 1 kHz and 10 Hz. Significant tests were conducted on the normalised RMS data in this figure (p < 0.0001).

Introducing 1 mM ascorbic acid solution to the trapped ferritin led to a 21% increase in RMS, while a concentration of 5 mM resulted in a 110% increase, indicating an accelerated rate of iron chelation and subsequent release of Fe^2+^ ions. At 10 mM, the RMS value remained higher than that of 1 mM but lower than 5 mM, exhibiting slower fluctuation in transmission signals with larger amplitude as indicated by its PDF in Figure 2d. This is attributed to the occurrence of saturation kinetics, whereby the available interactive sites between ascorbate and iron core become limited to generate surface complexes, which aligns with other ferritin work.^37^ The whole trace of ascorbic acid loading and the repeated results confirm that the high ascorbic acid concentration leads to increased dynamic motion of ferritin (Figures S2 and S3).

At high concentrations of ascorbic acid (1.5 M) when the solution becomes strongly acidic, we observed that ferritin underwent disassembly (Figure S4a). It takes approximately 12 minutes for the ferritin to be fully disassembled and escape from the trap in a stepwise fashion. During this time, the RMS value increased due to the unfolding of the ferritin channels (Figure S4b-d).^23,38^ Moreover, the presence of 1.5 M ascorbic acid resulted in a pH value of around 2, creating an acidic environment that further contributed to the disassembly of ferritin. To identify the impact of an acidic environment on ferritin’s structure, we focused on the conformational dynamics of single ferritins subjected to varying pH conditions *in situ*.

### Tracking disassembly kinetic and fragmentation of single ferritins

To understand the effect of pH on ferritin dynamics, we trapped single ferritin in pH 7.0 and replaced its environmental solution with acidic pH sequentially from 6.0 to 2.0 (see transmission traces in Figure 3a). The normalised RMS of the traces at pH 7.0 to pH 3.0 are relatively stable with values between 0.005 to 0.01 (Figure 3b), suggesting a stable structure within these solutions. Ferritin comprises 24 subunits categorised as heavy (H) or light (L), with the L-chain being more stable due to hydrogen bonding and salt bridges among subunits.^12,40^ The equine spleen ferritin used in this work has a relatively stable structure at acidic pH due to the dominant L-chain subunits.^2,12,40^ However, upon changing the pH to 2.0, the RMS jumps by 140%, indicating that the protein becomes highly dynamic. The mechanical instability of ferritin at pH 2.0 is attributed to protein swelling, monomer rotation, and subsequent dimer movement within the 3-fold channel, leading to channel opening. ^12,23,41^

**Figure 3.**
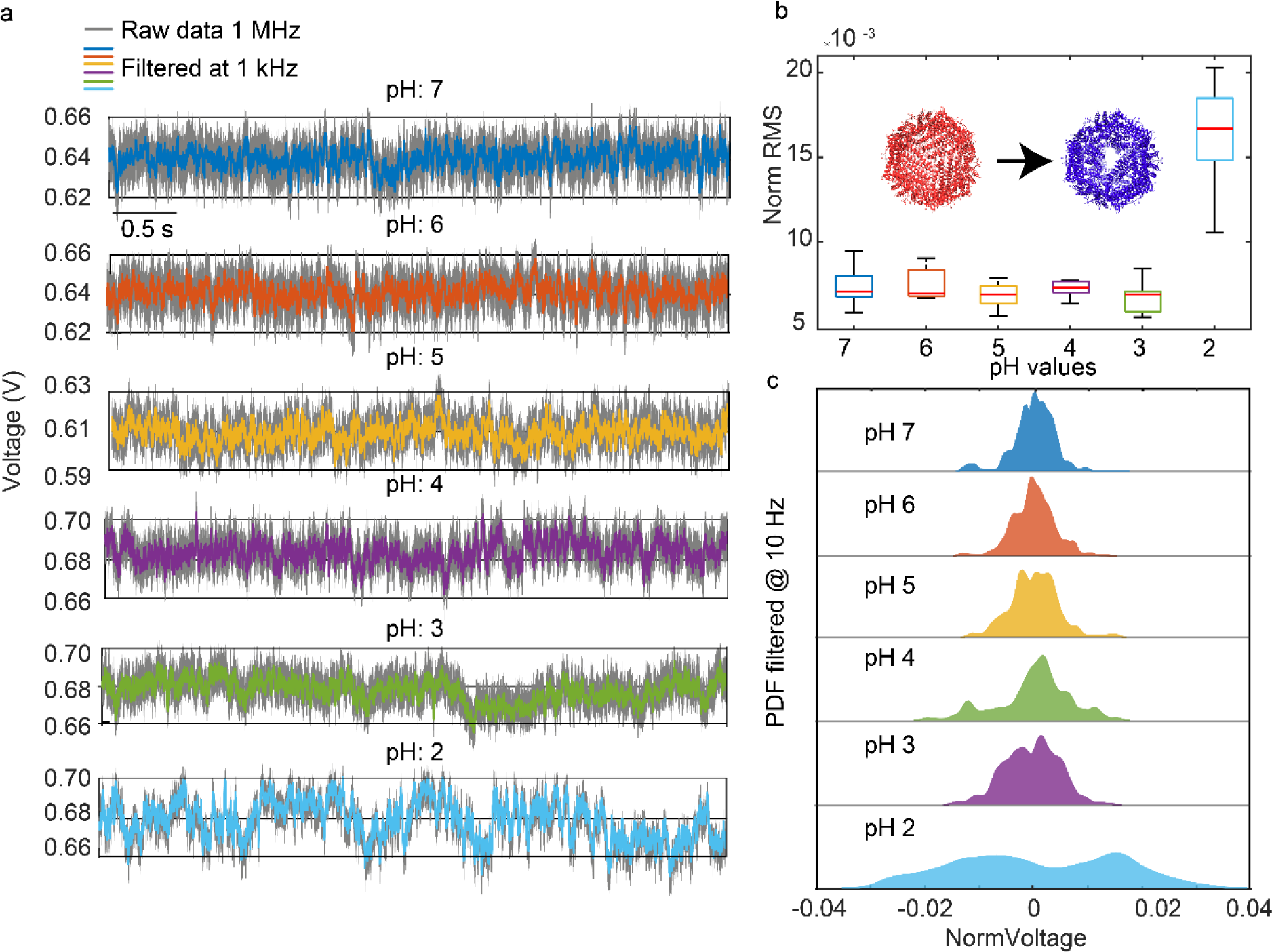
Conformational dynamics of trapped single ferritin exposed to solutions with different pH values. (a) Five-second transmission signals of the same single ferritin exposed to solutions with pH values of 7.0, 6.0, 5.0, 4.0, 3.0, and 2.0 in sequence (b) Normalised RMS of the transmission signal for a single ferritin exposed to pH values. And cartoon shows the opening of the 3-fold channels upon introducing to acidic pH 2.0. The data were Gaussian-filtered at 1 kHz, and the box plot displays the 25th and 75th percentiles of the data (c) PDF of five-second traces corresponding to the transmission signal of single ferritin exposed to different pH values (filtered at 10 Hz). Data were acquired at 1 MHz. Significant tests were conducted on the Norm RMS data. The p-values for comparisons between all pH values, except pH 2, ranged from 0.002 to 0.8 (0.002 <p < 0.8). Additionally, the p-value for comparisons between pH 2 and the other pH values was less than 0.0001 (p <0.0001).

When exposed to the acidic solution for a long duration, ferritin subunits began to dissociate, causing protein disassembly. Figure 4a demonstrates the capability of our approach to track the full disassembly kinetic of ferritin after the chamber is filled with pH 2.0. The stepwise reductions in the transmission signal (Figure 4b) show the step-by-step disassociation of ferritin’s subunits. As previously reported,^27^ we expect a linear relationship between the particle volume (sphere) and the transmitted signal of DNH structures. This relationship allows us to follow the ferritin disassembly process quantitatively.

**Figure 4.**
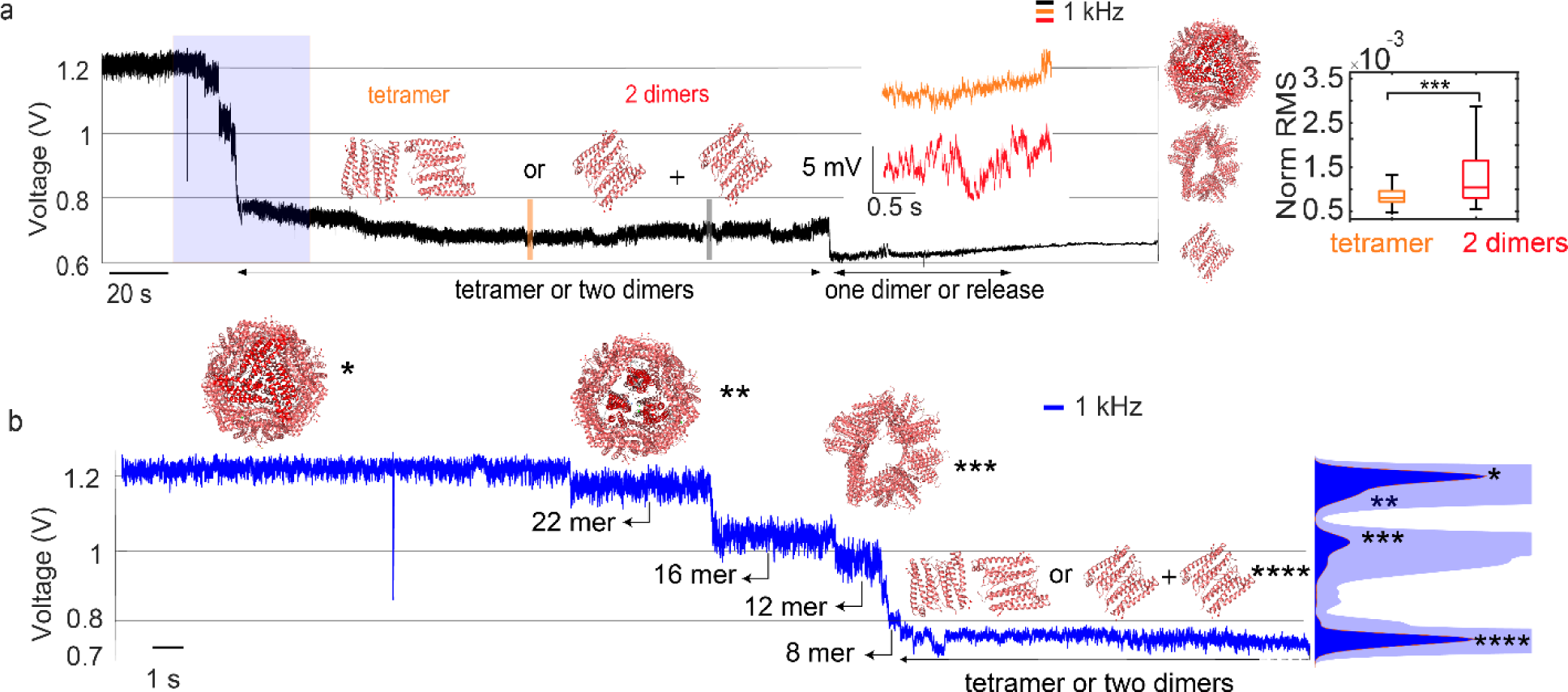
Disassembly process and kinetics of a single ferritin molecule into its subunits, as observed through changes in the transmission signal. (a) Full disassembly trace of a single ferritin with multiple significant drops in the transmission signal upon ferritin exposure to pH 2.0. Co-presenting of two fragments (two dimers) or one tetramer in the DNH hotspot along with their corresponding normalised RMS (right panel). The crystal structures on the right (PDB: 1IER) represent the structures corresponding to the transmission levels (b) An enlarged trace from panel a (highlighted part) focusing on the fragmentation process of ferritin. The asterisks mark the levels corresponding to the subunits of 20-mer, 16-mer, 12-mer, 8-mer, and tetramer or two dimers, matching with the peaks of the PDF on the right (with the transparent blue as the magnified PDF).

When ferritin disassembles into subunits, the molecular weight of the protein remaining in the trap reduces which consequently affects the transmission signal. The maximum transmission (*V*24mer) during the disassembly process represents the signal from a fully assembled ferritin with 24-mer subunits, and the transmission level at which the protein is released was labelled as *V*_DNH_. We linearly normalised the APD signal by taking the signal of trapped ferritin (*V*24mer) to be 100% and the signal of empty DNH (*V*DNH) to be 0%. By taking the sphere approximation, each subunit contributes to 4.17% of the optical signal. With this assumption, we correlated each drop of APD signal to the dissociation of a certain number of subunits.

Table 1 lists the obtained mean values of APD signals of each level in Figure 4, their normalised value, and the estimated number of subunits. We first observed a drop of 10%, corresponding to the dissociation of a dimer, leaving the ferritin to be a 22-mer subunit. Following this, the disassembly procedure continues by reduction of the trapping signal by 30, 45, 70, 80 and 95-100% which is attributed to the 16-mer, 12-mer, 8-mer, tetramer, and dimer respectively.

**Table 1.** Disassembly of single ferritin upon exposure to the pH 2.0.

| $V_{mean}$ (mV) | $\frac{V_{Xmer} - V_{DNH}}{V_{24mer} - V_{DNH}}$ | Ferritin<br>Fragments |
| --- | --- | --- |
| 1200 | 100% | 24-mer |
| 1154 | 91% | 22-mer |
| 1024 | 68% | 16-mer |
| 943 | 53% | 12-mer |
| 810 | 29% | 8-mer |
| 730 | 16% | 4-mer |
| 611 | 7% | 2-mer |
| 645 | 0% | released |

We note that the subunits always disassociate in single or multiple dimers, due to the well-known stable 2-fold axis of the ferritin. The 24 subunits of ferritin connect through the hydrogen bonding and electrostatic interactions, forming 2-, 3-, and 4-fold symmetries.^12,42,43^ Among these symmetries, the two-fold axis is the most stable one due to its largest number of interaction sites. Conversely, the three-fold axis is the weakest one because of its fewest interaction sites. ^12,43^ Therefore, there is a large chance that the initial disassembly of ferritin occurs in one of the three-fold channels.^12^

Among 4 disassembly experiments, except one experiment where the ferritin was released from the trap before the disassembly (Figure S5), three experiments exhibited a step-by-step disassembly signal. These experiments are denoted as Tests 1, 2, and 3, where Test 1 corresponds to the findings depicted in Figure 4 and Table 1, and the disassembly traces of Tests 2 and 3 can be found in Figures S6 and S7 respectively. The whole trace of protein during trapping and when exposed to the varying acidic pH conditions, and the disassembly signal are illustrated in Figure S8. All three transmission traces exhibit distinct drops after subjecting ferritin to pH 2.0 followed by protein release as the signal decreases to the baseline level. Figure 5a identified the number of remaining subunits based on the transmission level during the disassembly process, as described by Eq. 5. Where the *VXmer* is the mean transmission level of each step during disassembly.

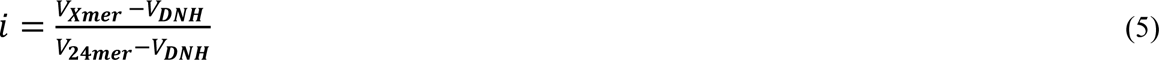

**Figure 5.**
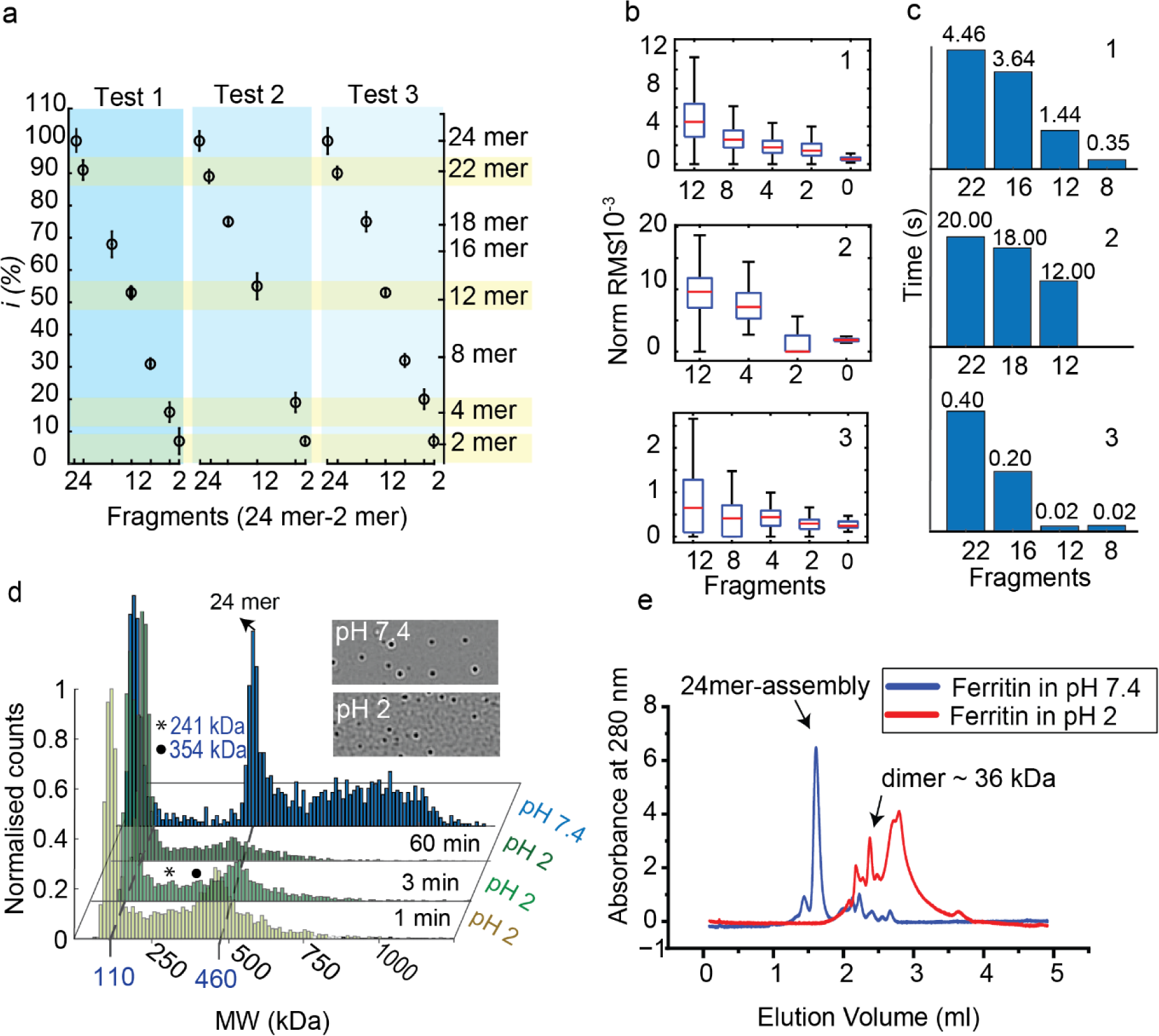
Single ferritin disassembly and its kinetics in three tests. (a) Number of subunits remaining in the trap during disassembly. (b) Normalised RMS value of disassembly steps from 12-mer to protein release in the three tests. (c) Dwelling time of the subunits in the trap in the three tests (tetramer and dimer dwelling times are excluded as tetramer can disassociate into two dimmers and still exist in the trap). (d) Mass photometry of ferritin at pH 2 across various time points (1 min, 3 min, and 60 min), along with ferritin at pH 7.4. The PDF is normalised to maximum counts for fair comparison among different measurements. The asterisk and circles mark the disappearance of intermediate subunits as ferritin is exposed to the acidic environment. Sample interferometric contrast images of binding events acquired from ferritin in pH 7.4 and pH 2. All images are on the same contrast scale with a field of view of 10.8 × 6.8 μm. (e) Size-exclusion chromatography of ferritin in pH 7.4 (blue) and pH 2 (red), see peak positions of protein markers in Figure S9.

The high sensitivity of optical nanotweezers enables us to discover a detailed pathway of the ferritin disassembly through different fragments (Figure 5a), which is otherwise difficult for other single-molecule techniques. We found that the disassembly of ferritin always starts with the disassociation of a dimer, forming a 22-mer assembly structure within the trap. This 22-mer fragment can go through various fragments during the disassembly. Despite the difference in the fragmentation pathway across the three tests, certain subunits (22-mer, 12-mer, 4-mer, 2-mer) consistently maintain their integrity across all three tests, as highlighted in Figure 5a. To the best of our knowledge, no other techniques can provide such detailed insights into ferritin disassembly.

The normalised RMS values presented in Figure 5b, spanning from 12-meric structures to the fragments released, agree with prior research that the RMS value of the transmission trace decreases with the size of the trapped globular proteins.^44^ To our surprise, the smaller fragment leaves the trap immediately after disassembly occurs. This fact is evidenced by the consistent RMS within each transmission level – if two fragments were coexisting inside the trapping site, their Brownian motion would lead to increased noise in the transmission trace compared to that of a single entity. We did not observe the coexistence of two fragments with the trapping site, except in the case of tetramer (or two dimers), where the fragments have identical size hence leading to significant noise difference, as shown in the transmission trace in Figure 4a and Fig. S7c.

The high RMS of the 12-mer fragment arises from its large accessible areas at the subunit-subunit interfaces, leading to the reduced number of hydrogen bonds between dimer-dimer interactions.^41^ We hypothesis that the iron core inside ferritin dissolves rapidly at this stage and makes the 12- mer fragment less stable.

Previous research on the disassembly and assembly kinetics of ferritin has employed techniques like circular dichroism with SAXS,^45,46^ but these methods primarily focused on the assembly process of ferritin in bulk measurement. Disassembly kinetics at the single-molecule level were not thoroughly investigated. The optical nanotweezers allow to resolve the dwell time of each subunit during disassembly down to milliseconds (Figure 5c). We observed a cooperative disassembly, where the disassembly of each subunit facilitates and accelerates the overall disassembly of the entire structure. This cooperative disassembly process has been demonstrated by simulation previously.^47^ In the case of tetramer and dimer, the dwell time might not be accurate as tetramers can dissociate into two dimers and these two fragments can co-exist in the trap as shown in Figure 4a.

We further validate our findings on single ferritin fragmentation at pH 2 using mass photometry as a complementary single-molecule approach, as demonstrated in Figure 5d. At pH 7.4 ferritin existed mainly at approximately 460 kDa, corresponding to its 24-mer full assembly structure. We also observed small fragments at 60 kDa, possibly due to the impurity of the protein sample, which is consistent with the Fast Protein Liquid Chromatography (FPLC) result in Figure 5e (blue). At pH 2, the dominant peak shifted to a lower mass (∼110 kDa). We identified two intermediate masses in the ferritin sample at pH 2, potentially corresponding to 12-mer (asterisk) and 18-mer (circle). These intermediate fragments decrease with prolonged exposure of ferritin to pH 2, reinsuring our observation that ferritin undergoes stepwise disassembling process. Although MP allowed us to comprehend the impact of acidic pH on the mass of ferritin fragments, it cannot provide details about their dynamic behaviour and their dwelling time during the disassembly of each subunit. Additionally, at lower molecular weights, as depicted in Figure 5e, it did not reveal the tetrameric and dimeric states of ferritin due to the intrinsic technical limit of the instrument when operating at more extreme conditions like low pH. However, by using FPLC we could confirm the dimeric state of ferritin at pH 2 (36 kDa), as it overlays well with the position of pepsin (Figure S9b), which is stable at pH 2 and has a molecular weight of 35 kDa. For further details about MP and FPLC in this work, see Supplementary Information Figures S9 and S10.

## Conclusion

This study used the DNH structures in the optical tweezer system to investigate the impact of different concentrations of ascorbic acid and acidic pH on single ferritins *in situ*. At the single-molecule level, we observed that ferritin’s dynamic gradually increases with the ascorbic acid concentration before reaching the saturation concentration (5 mM), suggesting a higher rate of Fe^3+^ to Fe^2+^ reduction from its core. When exposed to a high acid environment, *i.e.,* pH 2, ferritin became highly dynamic due to the protein swelling and hydrogen bonding protonation and eventually underwent a disassembly process. We identified the number of subunits during its disassembly pathway and discovered the most essential fragments consistently existing during the disassembly, which are 22-mer, 12-mer, 4-mer, and 2-mer. The 12-mer fragment is particularly significant, as it shows the highest protein dynamics of all subunits. This study further provides the kinetic of single ferritin disassembly under acidic pH conditions, revealing cooperative disassembly kinetics. Overall, these findings on the single-molecule level provide valuable insights into the potential engineering of ferritin’s structure to achieve desired functions like designing molecular machines and drug delivery platforms.^48,49^ Furthermore, this study showcases the quantitative analytical potential of optical nanotweezers in biomolecule analysis. This capability extends to diverse protein applications in future research and lays the groundwork for other single-molecule techniques, including solid-state nanopores, and AFM in investigating ferritin.^50,51^

## Method

### Fabrication of DNH structure

The double-nanohole (DNH) structures used in this study were fabricated by following the method outlined in the previous works.^25,27^ Initially, 550 µm thick fused silica wafers were coated with a 30-nm silicon nitride (SiNx) layer using low-pressure chemical vapour deposition (LPCVD) at 800°C. Subsequently, a 5-nm adhesion layer of titanium (Ti) and a 100-nm layer of gold (Au) were deposited on the silicon nitride using electron beam evaporation (Leybold Optics LAB 600H) at 190°C. The silica wafers were then diced into 10 mm × 10 mm chips using a dicing machine (Disco DAD321).

To create the DNH structures within the gold film, a focused ion beam (FIB) with a gallium ion source (Zeiss Crossbeam) was employed. The DNH geometry consisted of two circles with a diameter of 160 nm, spaced 200 nm apart, and connected by a 3 nm × 40 nm rectangle. The FIB parameters were optimised with an ion beam energy of 30 kV, a beam current of 1 pA, and dwelling times of 1.25 μs for the circles and 5 μs for the rectangle. These parameters ensured to create DNH structures with gap sizes predominantly around 20 nm, devoid of any residual gold. The scanning electron microscopy (SEM) image of the DNH at a 20° tilt is depicted in Figure S1.

### Optical nanotweezer setup

The optical components were obtained from Thorlabs, as previously described.^25^ The laser was focused to a spot of approximately 1.2 μm diameter using a 60× Plan Fluor objective (Nikon, Tokyo, Japan) with a numerical aperture (NA) of 0.85. A half-wave plate was used to adjust the laser’s polarisation to be perpendicular to the line connecting the centres of two holes. The power density at the DNH sample was 19 mW/µm^2^, resulted from the incident laser power of 32 mW before the objective. The transmitted light was collected by a silicon avalanche photodiode (APD120A, Thorlabs), which converted the optical intensity into a voltage signal. The APD’s voltage signal was recorded by a data acquisition card (USB-6361, NI) at a sampling rate of 1 MHz using a customised LabVIEW program.

### Preparation of proteins, different pH buffers, and ascorbic acid solutions

#### Protein solutions

We procured holo-ferritin (derived from equine spleen, F4503), along with other chemicals, from Sigma-Aldrich in the United Kingdom. For the trapping experiments, we employed 0.5 μM of ferritin in 0.1 M phosphate buffer (PB) with a pH value of 7.4.

#### Different pH solutions

To create solutions with varying pH levels, we adjusted the pH by adding hydrochloric acid (HCl) with a concentration of 0.1M (purchased from Fisher Scientific, product code 15697310) to deionized water. In addition to testing the pH change in deionised water, we conducted experiments in both pH 7.4 phosphate buffer (PB) and a Glycine-HCl buffer 0.1 M at pH 2.0 The Glycine-HCl buffer served as a control to assess the effects of a sudden pH shift from 7.4 to 2.0.

#### Ascorbic acid-buffer solution

L-Ascorbic acid (A5960) was dissolved in PB buffer (pH 7.4) at three different concentrations (1 mM, 5 mM, and 10 mM) freshly before each test.

### Fluidic system

The flow cells used in this study are identical to those previously reported.^25,27^ We produced the flow cells using a FormLab 2 printer, employing Clear V4 resin with a resolution of 50 µm (Formlabs Inc., USA). Samples in the flow cell were sealed with a cover glass using a two-component silicone glue (Twinsil, Picodent, Germany). The DNH sample and cover slide were separated by a 50 µm thick double-sided tape (ARcare92712, Adhesive Research, Inc.), creating a fluidic channel with a volume of 3.5 µL. A syringe pump (Harvard Apparatus, US) controlled the flow rate and direction through a 12-port valve (Mux Distrib, Elve Flow, France). To replace the buffer after protein trapping, we introduced buffer into the flow chamber at a flow rate of 3 µL/min. Considering the internal diameter of the tubing in the flow controller system and mentioned flow rate, the different pH and ascorbic acid solutions reach the flow chamber 25 µL (6 minutes after injection) after passing through the flow controller.

### Size exclusion chromatography

The pH-induced disassembly of ferritin was investigated through size exclusion chromatography (SEC) utilising a Superose 6 Increase 3.2/300 column. Molecular sizes of both the intact ferritin assembly and its constituent fragments were determined under various conditions by comparing their elution volumes to those of standard proteins, namely Aprotinin (6.500 kDa, Sigma-Aldrich A3886), Carbonic Anhydrase (29 kDa, Sigma-Aldrich C7025), Bovine Serum Albumin (66 kDa, Sigma-Aldrich A8531), Alcohol Dehydrogenase (150 kDa, Sigma-Aldrich A8656), Beta-amylase (200 kDa, Sigma-Aldrich A8781), and Thyroglobulin (669 kDa, Sigma-Aldrich T9145). The concentration of all protein markers was maintained at 0.165 mg/ml, except for ferritin, which was at a concentration of 1.65 mg/ml.

To assess retention times and ensure the accurate positioning of the peak at pH 2, pepsin, a stable protein under acidic conditions with a molecular weight of 35 kDa, was used at a concentration of 0.4 mg/mL. The mobile phase composition consisted of 10 mM PB buffer with 50 mM sodium chloride, pH 7.4, and for acidic conditions, 100 mM Glycine-HCl at pH 2. All SEC experiments were conducted using the ÄKTA-Explorer FPLC system with protein detection by absorption at 280 nm.

### Single-molecule mass photometry analysis

MP measurements were performed using a TwoMP mass photometer (Refeyn), adhering to the manufacturer’s calibration guidelines. Native Mark^TM^ (Thermo Fisher Scientific) was employed for calibration in T50 buffer at pH 7.4. The calibration accuracy was estimated to be 1.5% (R2=0.99997). The experiments were carried out using conventional microscope cover glasses (Marienfeld, no 1.5 H) cleaned by rinsing with deionized water (X5) and isopropanol (X5) followed by drying under a N2 flow, using a silicon gasket (Grace Bio-labs) to confine the sample. For the entire measurement process, a total volume of 20 μL of mixed ferritin in buffers (50 nM) was used, which was then introduced into the pristine wells of the gaskets. To ensure procedural cleanliness, MP measurement buffers underwent filtration using 0.22 μm syringe filters. The protein adsorption events were obtained using AcquireMP software (Refeyn) after recording 60s videos for each sample. Data analysis was performed using DiscoverMP software (Refeyn) and OriginPro 2021 (OriginLab).

### Data analysis

In this study, MATLAB scripts were employed for the analysis of all presented data. Raw data underwent filtration using a zero-phase Gaussian low-pass filter with a specified cut-off frequency (1 kHz or 5 kHz) applying the filtfilt.m function. Probability density functions (PDF) were determined using the ksdensity.m function and filtered at 10 Hz. The normalised root mean square (RMS) values (Figures 2, 3) were calculated by dividing the standard deviation of a 0.5-second trace by its mean value. The trance length for the last figure was 0.015-second due to the short duration of each step.

## Supporting Information

S1 - Scanning electron microscopy (SEM) image from the DNH sample; S2 - Optical properties of the DNH sample; S3 - Continuous trace of single ferritin across varying ascorbic acid concentrations, S4 - Repeating test of ascorbic acid’s effect on single ferritin’s dynamic at various concentrations, S5 - Effect of high concentration of ascorbic acid (1.5 M) on the single ferritin. S6 - Non-stepwise disassembly of single ferritin upon exposure to pH 2. S7 - Stepwise disassembly trace of test 2. S8 - Stepwise disassembly trace of test 3. S9 - Continuous trace of single ferritin exposed to varying pH values. S10 - Size exclusion chromatography (protein markers and pepsin). S11- Mass photometry

## Author Information Corresponding Authors

**Cuifeng Ying**- Advanced Optics and Photonics Laboratory, Department of Engineering, School of Science and Technology, Nottingham Trent University, Nottingham NG118NS, United Kingdom.

**Mohsen Rahmani-** Advanced Optics and Photonics Laboratory, Department of Engineering, School of Science and Technology, Nottingham Trent University, Nottingham NG118NS, United Kingdom.

## Authors

**Arman Yousefi-** Advanced Optics and Photonics Laboratory, Department of Engineering, School of Science and Technology, Nottingham Trent University, Nottingham NG118NS, United Kingdom

**Ze Zheng-** Advanced Optics and Photonics Laboratory, Department of Engineering, School of Science and Technology, Nottingham Trent University, Nottingham NG118NS, United Kingdom

**Saaman Zargarbashi-** Advanced Optics and Photonics Laboratory, Department of Engineering, School of Science and Technology, Nottingham Trent University, Nottingham NG118NS, United Kingdom

**Mahya Assadipapari-** Advanced Optics and Photonics Laboratory, Department of Engineering, School of Science and Technology, Nottingham Trent University, Nottingham NG118NS, United Kingdom

**Graham J. Hickman-** Interdisciplinary Biomedical Research Centre, School of Science and Technology, Nottingham Trent University, Nottingham NG118NS, United Kingdom

**Christopher D. J. Parmenter-** Nanoscale and Microscale Research Centre, University of Nottingham, Nottingham NG7 2RD, United Kingdom

**Carlos J. Bueno-Alejo-** School of Chemistry, University of Leicester, University Road, Leicester LE1 7RH, United Kingdom

**Gabriel Sanderson-** Advanced Optics and Photonics Laboratory, Department of Engineering, School of Science and Technology, Nottingham Trent University, Nottingham NG118NS, United Kingdom

**Dominic Craske-** Imaging Suite, School of Science and Technology, Nottingham Trent University, Nottingham NG118NS, United Kingdom

**Lei Xu-** Advanced Optics and Photonics Laboratory, Department of Engineering, School of Science and Technology, Nottingham Trent University, Nottingham NG118NS, United Kingdom

## Acknowledgments

We express our sincere thanks to Prof. Andrew J. Hudson for providing access to mass photometry at the University of Leicester. This research work has been kindly supported by the Royal Society and the Wolfson Foundation. The authors appreciate the use of the NTU Imaging Suite, NTU High-Performance Computing Cluster Hamilton, NTU Medical Technologies Innovation Facility (MTIF), and NTU Interdisciplinary Biomedical Research Centre (IBRC) - as well as the Nanoscale and Microscale Research Centre (nmRC). Strategic Longer and Larger Grant: Frontier Bioscience from the Biotechnology and Biological Sciences Research Council (How do RNA-binding proteins control splice site selection? BB/T000627/1)

## Notes

### Competing Interest Statement

The authors have declared no competing interest.

### Summary of Updates

New data from FPLC and Mass Photometry added.

