## Supplementary Information for "Structural Flexibility and Disassembly Kinetics of Single Ferritins Using Optical Nanotweezers"

### S1- SEM image of the double nanohole (DNH).

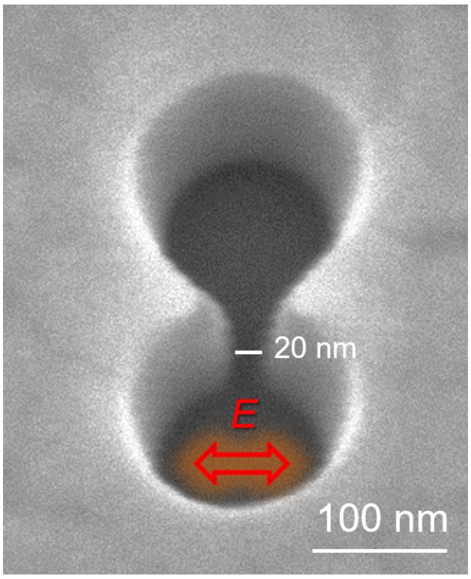


Figure S1. Scanning electron microscopy (SEM) image of the DNH structure with 20° tilted angle.

### S2: Optical properties of the DNH sample

The trapping mechanism in this system relies on the self-induced back action (SIBA) trapping, which offers the advantage of not being strictly limited to a specific excitation wavelength.[1] Figure 1b in the main text shows that DNH provides a highly confined field enhancement in the gap, enabling a strong gradient force for efficient trapping of single proteins.

*Finite Element Simulation*: The field distribution of DNH was simulated using the finite element method (FEM) by COMSOL Multiphysics 6.0. We considered a material stack consisting of a 100 nm Au layer, 5 nm of Ti, and 30 nm of $SiN_x$ on a fused silica substrate, surrounded by water with a refractive index (RI) of 1.333. Based on the SEM image in Figure S1 we analysed the gap geometry which exhibits a trapezium shape. The gap size, corresponds to a trapezium with a smaller base of 20 nm, as illustrated in Figure 1b, y-z plane in the main text. To account for tapering in the DNH structures, we introduced an 'n side truncated pyramid' structure with an RI of 1.333, transitioning from ~20 nm gap width at the bottom to ~45 nm at the top of the Au layer.

### S3: Continuous trace of single ferritin across varying ascorbic acid concentrations

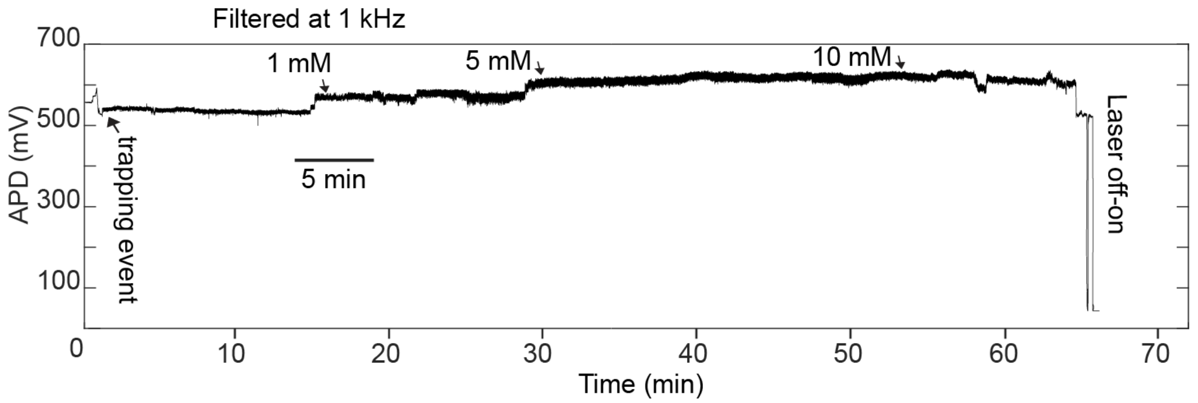


Figure S2. Continuous transmission signal of a single ferritin trapped in the hotspot of the DNH in PB buffer and exposed to the different concentrations of ascorbic acid (1 mM, 5 mM, and 10 mM). The transmission increased upon introducing 1mM and 5 mM concentrations of ascorbic acid to the ferritin. It is possible that the net charge of ferritin reduces in the acidic solution, weakening the electrostatic force and allowing it to approach the gold surface, resulting in a change in transmission. It is unlikely that these jumps correspond to the trapping of additional ferritins, as the fluidic chamber has been flushed with a solution that contains no ferritins.

### S4: Repeating test of ascorbic acid's effect on single ferritin’s dynamic at various concentrations

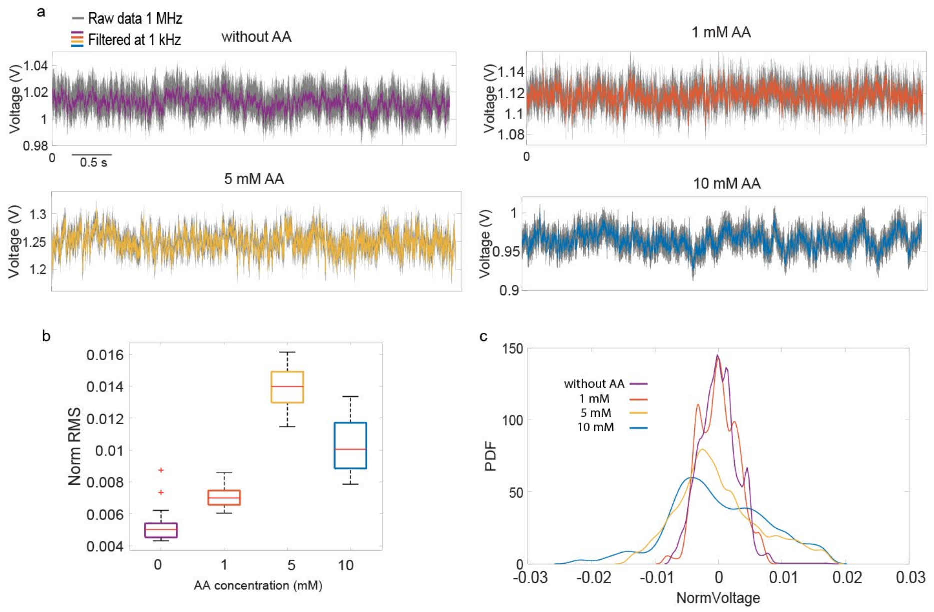


Figure S3. (a) Transmission signal of trapped ferritin at different concentrations of ascorbic acid (1 mM, 5 mM, and 10 mM). (b) Normalised root-mean-square (RMS) of 5 second transmission signal when ferritin is trapped in PB buffer and introduced to different ascorbic acid concentrations. The data were Gaussian-filtered at 1 kHz, and the box plot displays the 25th and 75th percentiles of the data (c) PDF of 5-seconds traces of single ferritin and different ascorbic acid concentrations (filtered at 10 Hz). Data were acquired at 1 MHz and then Gaussian filtered at 1 kHz.

### S5: Effect of high concentration of ascorbic acid (1.5 M) on the single ferritin.

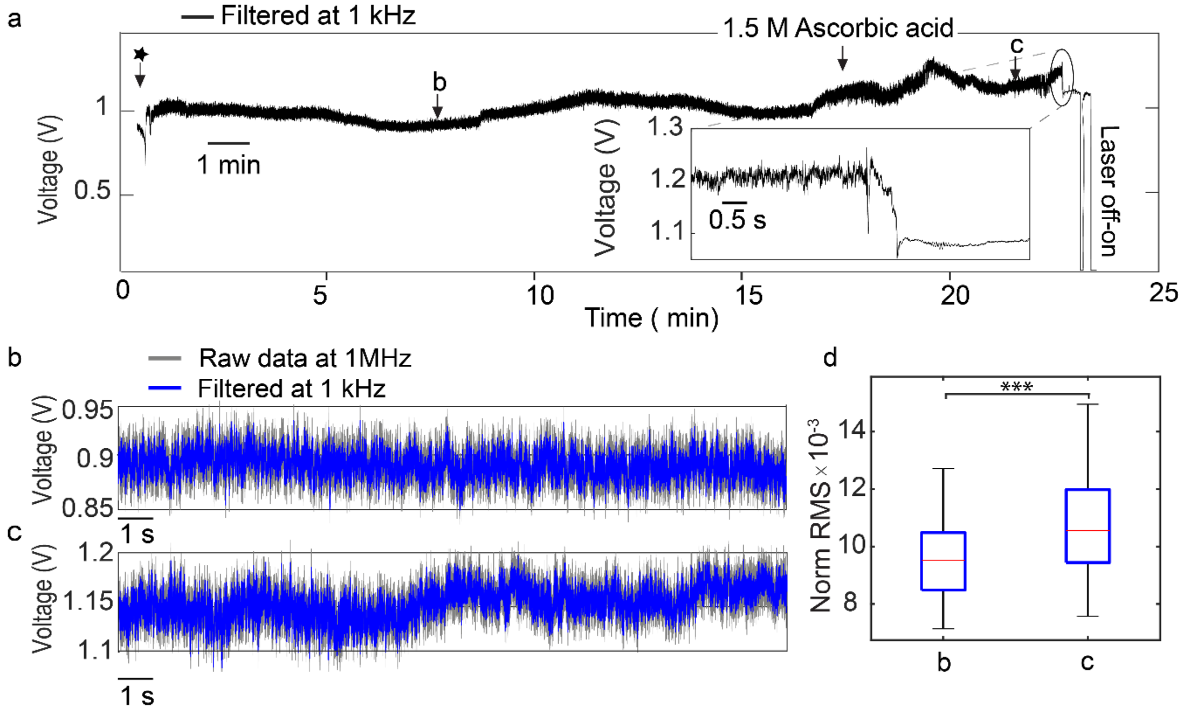


Figure S4. Disassembly of a single ferritin exposed to 1.5 M ascorbic acid. (a) 25 mins transmission trace depicts the protein trapping (indicated by the star), exposure to 1.5 M ascorbic acid (arrow), disassembly (magnified in the inset), and release by turning off the laser. (b) 20-s magnified transmission signal from panel a (marked by the arrow). (c) 20-s magnified transmission signal from panel a during exposure to 1.5 M ascorbic acid. (d) Boxplot of normalised RMS for 20-second transmission traces shown in panels b and c, collected at 0.1-second intervals at 1 kHz frequency. Asterisks indicate statistically significant differences (***$p < 0.0015$).

### S6: Non-stepwise disassembly of single ferritin upon exposure to pH 2.

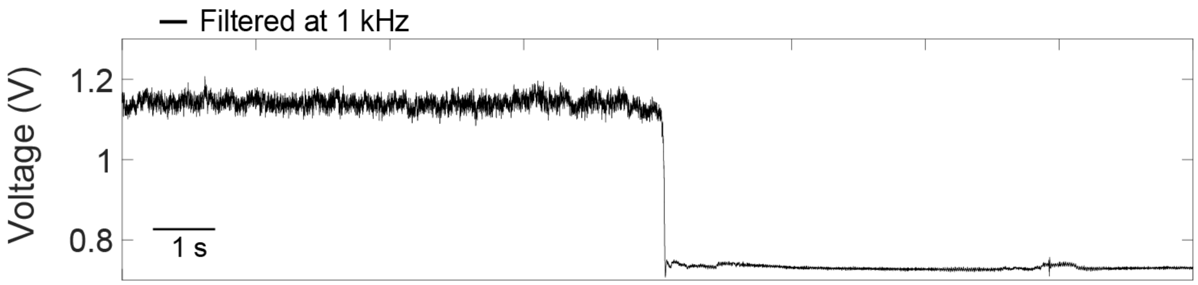


Figure S5: a trace of Single ferritin disassembly upon exposure to pH 2 showing non-stepwise behaviour. Data acquired in 1 MHz, in this figure, 1 kHz filtered data is showcased.

### S7: Stepwise disassembly trace of test 2

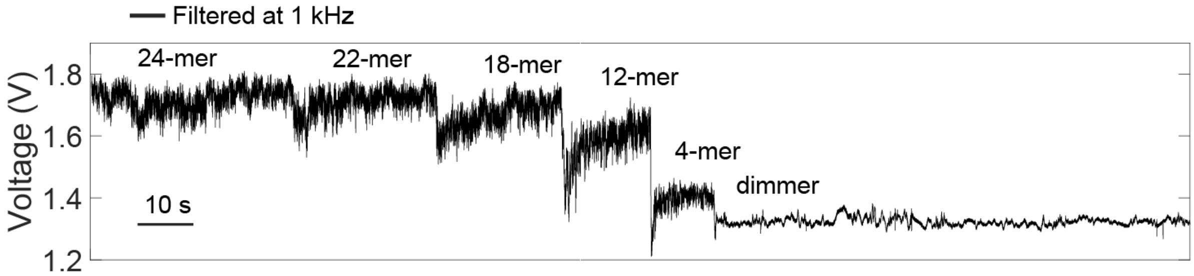


Figure S6. The whole traces of step-wise disassembly for test 2. Data was acquired in 1 M Hz, and filtered to 1 kHz.

### S8: Stepwise disassembly trace of test 3

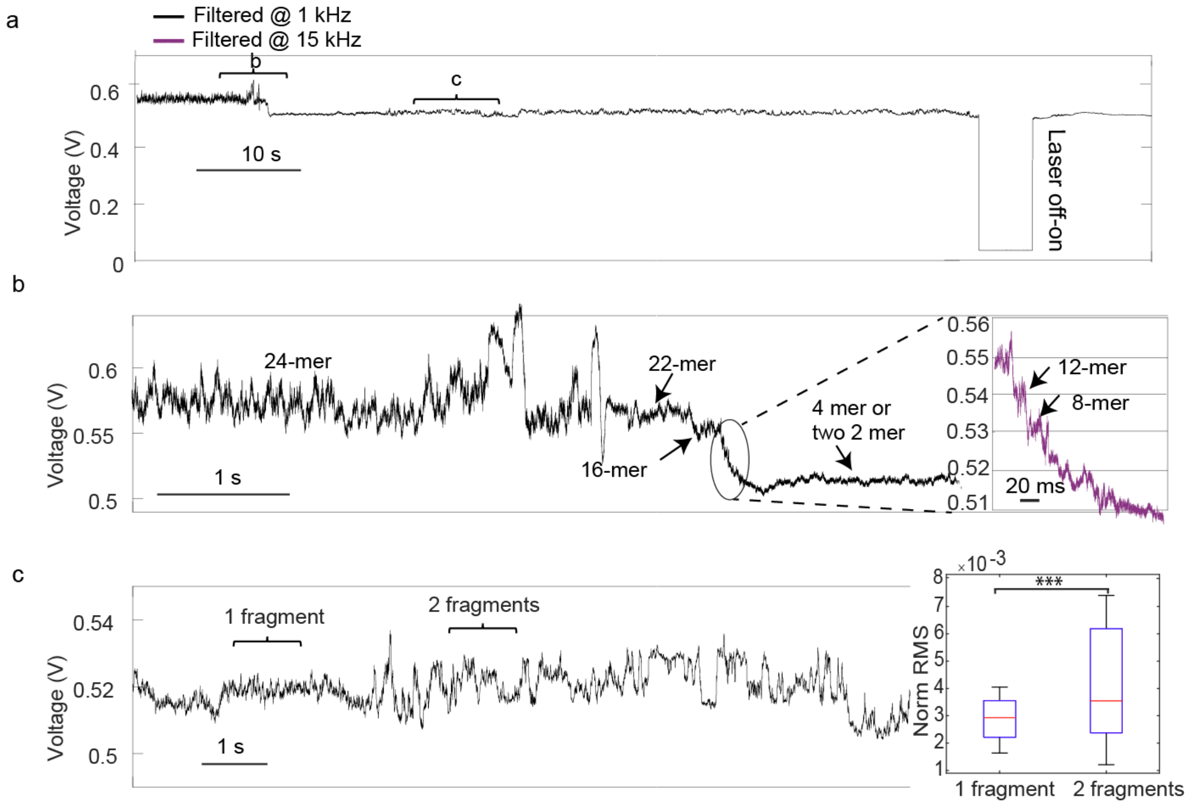


Figure S7 Disassembly trace of test 3. (a) Transmission trace showing the disassembly of ferritin at pH 2.0, encompassing of turning off the laser to release the dimer fragments (two fragments) from the trap. (b) A zoomed-in view of the ferritin disassembly, highlighting the evident stepwise progression. The right panel image is an enlarged trace section, emphasising the disassembly steps. (c) Enlarged section of the transmission signal when two dimer fragments are both in hotspot of the DNH. The right panel depicts the normalised RMS of the 2-second transmission segments marked in the trace, indicating the presence of one tetramer and two dimers in the trap.

**S9: Continuous trace of single ferritin exposed to varying pH values.**

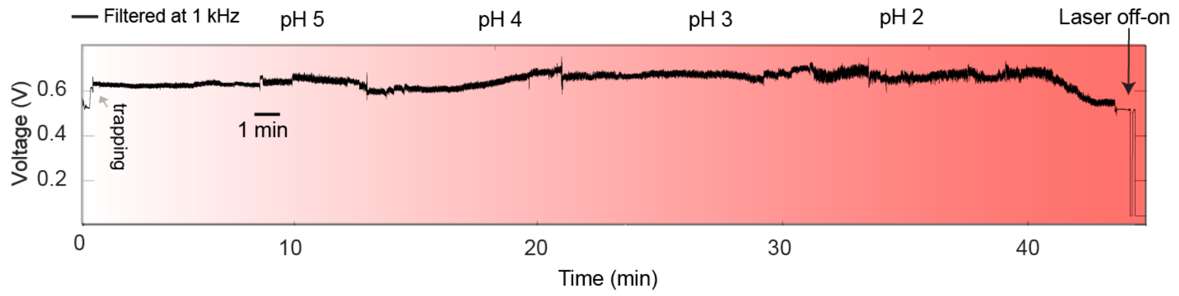


Figure S8. Continuous transmission trace of a DNH with a ferritin protein trapped and subsequently subjected to pHs of 5, 4, 3 and 2. After exposure to pH 2, the transmission signal decreased over time, indicating a protein disassembly process. The pH levels varied from 5 to 2. The intensity of the red hue increased proportionally with the decreasing pH, indicating a change in the difference in absolute pH values upon introduction to the chamber - turning off the laser for 5 seconds, and the transmission signal returned to the baseline, suggesting the fragments were released.

**S10: Size exclusion chromatography (protein markers and pepsin)**

To confirm the purity of ferritin at pH 7.4, ferritin solution was examined by fast protein liquid chromatography (FPLC) with various protein markers. Additionally, to pinpoint the dominant ferritin fragments at pH 2, we conducted FPLC with pepsin (35 kDa), chosen for its molecular weight proximity to the dimeric state of ferritin and its stability in an acidic environment.

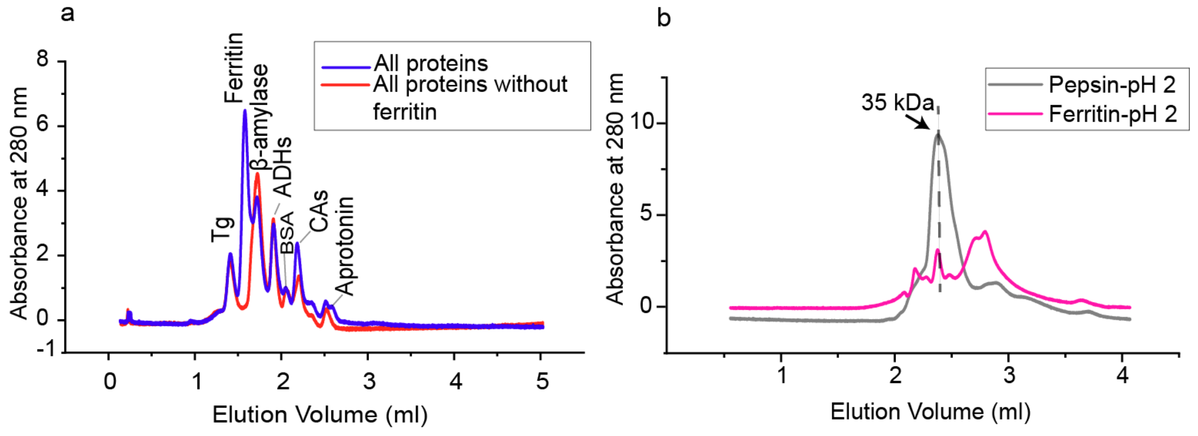


Figure S9. FPLC chromatograms of proteins. (a) marker proteins including ferritin (blue), and marker proteins excluding ferritin (red) to confirm the peak position of ferritin. (b) Chromatograms of pepsin and ferritin proteins in pH 2.

### S11: Mass photometry

Single-molecule mass photometry (SMMP) is a label-free technique that assesses the molecular mass of individual molecules, particularly biomolecules ranging from 40 kDa to 5 MDa within a sample. The alteration in reflectivity at the glass-water interface, induced by the binding of a biomolecule from the solution to the coverslip surface, leads to a localised change in contrast.[2] SMMP results in Figure S9 display peaks representing the native 24-mer assembled ferritin in T50 buffer at pH 7.4. Additionally, a broad peak of the 48-mer is evident in the figure, suggesting the potential binding of two ferritin molecules together. Furthermore, distinct fragments around 60 kDa, separate from the fully assembled ferritin, are evident in the figure.

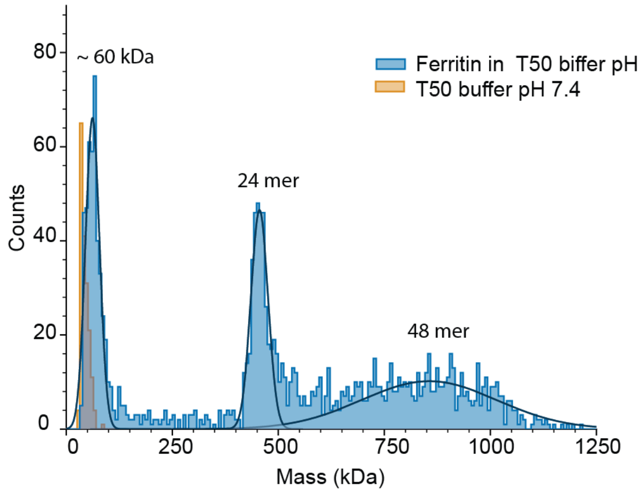


Figure S10. SMMP of T50 buffer pH 7.4 with and without 50 nM ferritin
